# In Silico Generation of Gene Expression profiles using Diffusion Models

**DOI:** 10.1101/2024.04.10.588825

**Authors:** Alice Lacan, Romain André, Michele Sebag, Blaise Hanczar

**Affiliations:** IBISC, University Paris-Saclay (Univ. Evry); TAU, CNRS-INRIA-LISN, University Paris-Saclay; Diagnostic Image Analysis Group, Department of Medical Imaging, Radboud University Medical Center

## Abstract

**Motivation:** RNA-seq data is used for precision medicine (e.g., cancer predictions), which benefits from deep learning approaches to analyze complex gene expression data. However, transcriptomics datasets often have few samples compared to deep learning standards. Synthetic data generation is thus being explored to address this data scarcity. So far, only deep generative models such as Variational Autoencoders (VAEs) and Generative Adversarial Networks (GANs) have been used for this aim. Considering the recent success of diffusion models (DM) in image generation, we propose the first generation pipeline that leverages the power of said diffusion models.

**Results:** This paper presents two state-of-the-art diffusion models (DDPM and DDIM) and achieves their adaptation in the transcriptomics field. DM-generated data of L1000 landmark genes show better predictive performance over TCGA and GTEx datasets. We also compare linear and nonlinear reconstruction methods to recover the complete transcriptome. Results show that such reconstruction methods can boost the performances of diffusion models, as well as VAEs and GANs. Overall, the extensive comparison of various generative models using data quality indicators shows that diffusion models perform best and second-best, making them promising synthetic transcriptomics generators.

**Availability and implementation:** Data processing and full code available at: https://forge.ibisc.univevry.fr/alacan/rna-diffusion.git

**Supplementary information:** Supplementary data are available at *BioRxiv* online.

## 1. Introduction

Precision medicine is advancing rapidly by leveraging multimodal patient data, ranging from medical imaging and electronic health records (EHRs) to genomics. The potential of each data source is investigated extensively, using deep learning in computer vision or natural language processing (NLP) for, respectively, medical imaging and EHRs, yet more modestly in genomics. The genomics domain, encompassing a variety of high-dimensional data like transcriptomics, holds valuable information regarding the prediction of patients’ diagnoses, prognoses, and treatment responses. Next-generation sequencing (NGS) methods (Hong *et al*., 2020) make it possible to afford significant RNA sequencing (RNA-seq) data at a lower cost and high accuracy. This data surge is set to fundamentally impact precision medicine to develop person-dependent treatments. For instance, bulk RNA is increasingly used to predict tissue-specific conditions, cancer diagnoses, and prognoses (Kourou *et al*. 2015; Katzman *et al*. 2018; Kim et al. 2020).

On the methodological side, advanced deep learning methods awaken great hopes as they enable representing and learning complex non-linear relationships from high-dimensional data (Jha *et al*., 2022). In practice, however, the current scarcity of gene expression data (Koumakis, 2020) remains a crippling problem for the use of deep learning. Deep models notoriously require massive amounts of training data, all the more so in high-dimensional spaces (circa 22,000 coding genes). In transcriptomics, the number of samples is restricted due to the limited number of patients, residual costs of RNA sequencing methods, and the need for cohort integration.

This scarcity makes for insufficiently representative datasets, resulting in a poor generalization, that is, overfitting. Building efficient deep neural models from small datasets thus often relies on domain knowledge and computationally intensive efforts (e.g., to regularize the learning criterion and adjust the method hyperparameters).

The automatic generation of sizeable synthetic transcriptomics datasets, faithful to the data distribution, is a sound way to address the shortage of data. The generation of synthetic samples can also enable the target tasks to be conducted on anonymized synthetic data, thus addressing privacy concerns (Yale *et al*., 2020). In computer vision and NLP, *data augmentation* is a regularization technique exploiting additional artificial training samples that follow the same data distribution as the original training set. Deep generative models like Variational Auto-Encoders (VAEs; Welling and Kingma, 2014) and Generative Adversarial Networks (GANs; Goodfellow *et al*., 2014) have been presented as good candidates for generating such synthetic samples (Shorten and Khoshgoftaar, 2019).

For cancer applications, a promising WGAN-GP data generation strategy was proposed by Viñas *et al*. (2021); the classifier trained on generated samples yielded state-of-the-art prediction results on bulk RNA data. Moreno-Barea et al. (2022), Li et al. (2023), and Lacan et al. (2023) presented GAN-based data augmentation approaches to boost performances using conditional GANs and better evaluate data quality. Interestingly enough, the latent representation of VAEs and GANs were also leveraged to capture biologically relevant features for cancer-related tasks (Way and Greene, 2018) and interpretations of biological pathways (Seninge *et al*., 2021). However, Lacan et al. (2023) showed that synthetic data diversity and mode collapse remained challenging for such generators.

Recently, a new type of generative model referred to as diffusion models (DMs) (Sohl-Dickstein *et al*., 2015) are revolutionizing image generation with, for example, DALL-E 2 (Ramesh *et al*., 2022). Diffusion models are celebrated for their ability to generate a large diversity of high-quality samples and offer new interpolation possibilities. While DMs have been successfully applied to protein generation with FoldingDiff (Wu *et al*., 2024), they have not yet been applied to transcriptomics (except for Wang *et al*., 2023, reporting an unsuccessful application). A primary challenge lies in the transcriptomics data’s high dimensionality and complex structure (e.g., involving long-range relations).

Building upon the methodology used to deploy and assess data augmentation with VAEs and GANs (Lacan *et al*., 2023), the proposed contribution is a gene expression generation pipeline leveraging the power of diffusion models. The critical issues related to the high dimensionality of transcriptomics data are handled by considering the so-called 1,000 landmark genes (Subramanian *et al*., 2017): the data generation is conducted in this reduced 1,000-dimension space, and the generated samples are mapped to the remaining genes space using a trained linear or non-linear approach (Chen *et al*., 2016; Jeon *et al*., 2022). To the best of our knowledge, this is the first success in applying the famed diffusion models on bulk RNA data.

## 2. Diffusion-based generation method

This Section presents the core contribution of the paper, the Diffusion Model-based data generation methodology illustrated in Fig.1. For the sake of containedness, the formal background of DMs is first introduced and discussed, referring the reader to the original papers for a more comprehensive presentation, and to Appendix J for details on the baseline VAEs and GANs. An original methodology is also proposed to address the high dimensionality issue inherent in transcriptomics data, which constitutes a significant hurdle for DM-based generative models.

### Notations

In the following, 𝒟_*t*_ denotes the (unknown) true data distribution, and 𝒟_*g*_ the learned distribution parameterized from the model parameters *θ*. Deep generative models proceed by learning a distribution 𝒟_*g*_= *p*_*θ*_(*x*) that approximates the target distribution 𝒟_*t*_, supporting a sampling mechanism.

### 2.1 Diffusion Models

Early success in stable diffusion-based generative modeling is due to Sohl-Dickstein et al. (2015). However, widespread interest arose with the introduction of Denoising Diffusion Probabilistic Models (DDPMs) by Ho *et al*. (2020) and Dhariwal and Nichol (2021), outperforming GANs in terms of class and mode coverage, image quality, and stability, making them prominent generative models.

### Denoising Diffusion Probabilistic Models

Ho *et al*. (2020) introduce a Markov Chain framework called diffusion. In forward mode, starting from an initial example *x*_0_ *~ 𝒟*_*t*_, a sequence of *T* samples *x*_1_, … *x*_*T*_ is simulated, where *x*_*t*_ is obtained by adding Gaussian noise to *x*_*t−*1_ (Fig. 1, panel B):

**Fig. 1:**
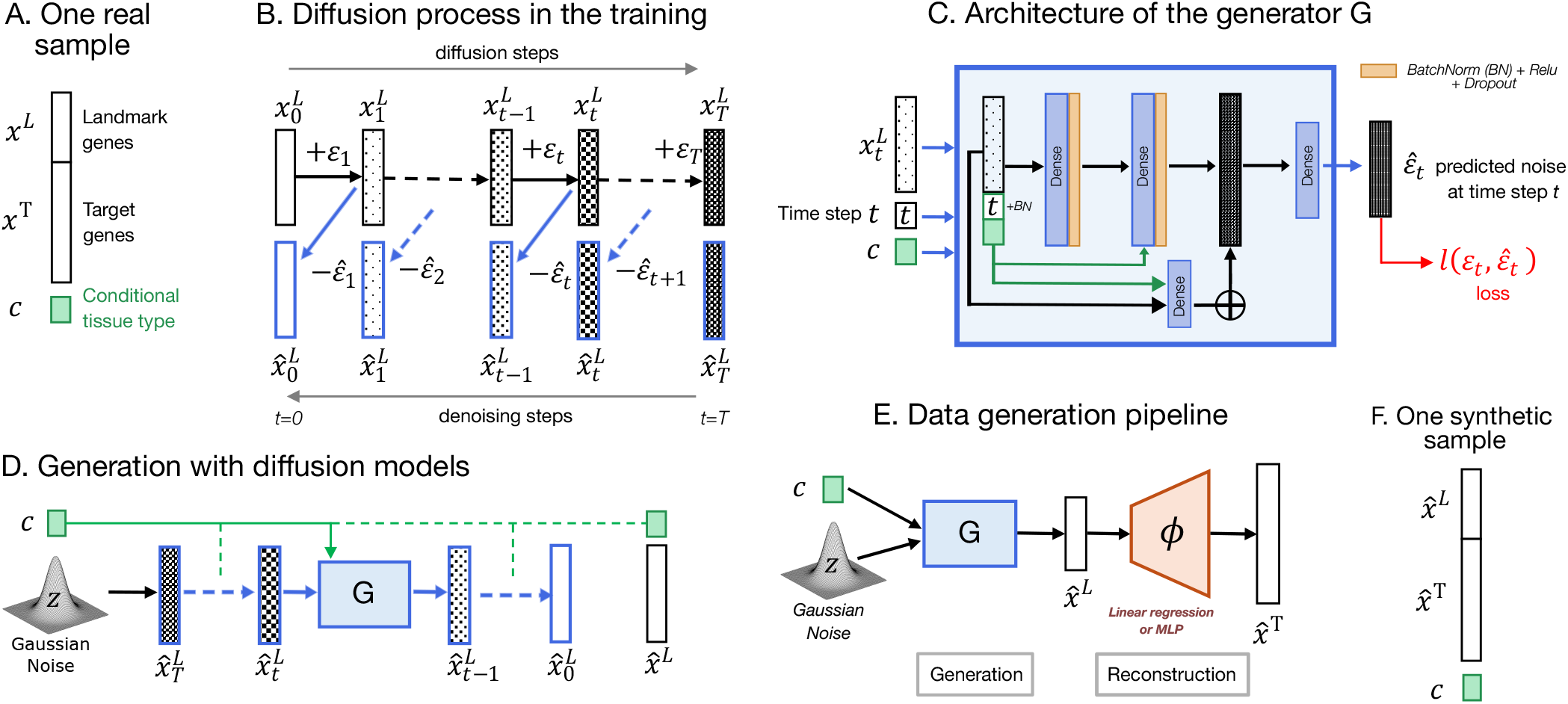
Overview of our data generation pipeline with a diffusion model (DM). For each real sample (A) we consider the L1000 landmark genes *x*^*L*^ and the remaining target genes *x*^*T*^, as well as a corresponding covariate *c* (tissue type). During the training of the DM (B), the model (C) learns to progressively denoise the samples for which gaussian noise *ϵ*_*t*_ was added at each time step *t*. The model takes as input a noisy sample 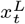 and directly predicts this added noise. It is conditioned on both the time step *t* and the tissue type *c*. For the generation process (D), the DM takes as input Gaussian noise and progressively denoises the sample, exactly like in training (B). The overall pipeline (E) consists of a generation phase, followed by a reconstruction phase taking as input the generated landmark genes 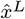 to recover the final synthetic sample with target genes 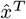 (F).

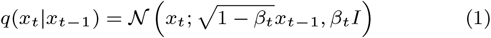

The noise level *β*_*t*_ ∈ [0, 1] gradually increases along a noise schedule, ensuring that the final *x*_*T*_ is essentially a pure Gaussian noise sample. A specifics of the DM forward process is that the intermediate *x*_*t*_ are in the same input space as *x*_0_: the forward process only depends on the (fixed) noise schedule *β*_*t*_.

Besides the iterative process (Eq. 1), the forward process can also be computed in closed form as in Eq. 2, enabling the direct modelling of all noisy *x*_*t*_ from the initial sample *x*_0_ and Gaussian noise *ϵ ~ 𝒩* (0, *I*_*d*_).

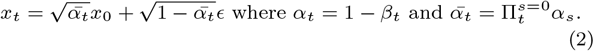

The closed-form computation speeds up training compared to the iterative computation; it promotes a stable setting and alleviates memory storage issues.

The generative model corresponds to the backward process trained to invert the forward process, achieving the progressive denoising of each sample *x*_*T*_ (Fig. 2, left to right). Formally, the generator learns the reverse process defined as:

**Fig. 2:**
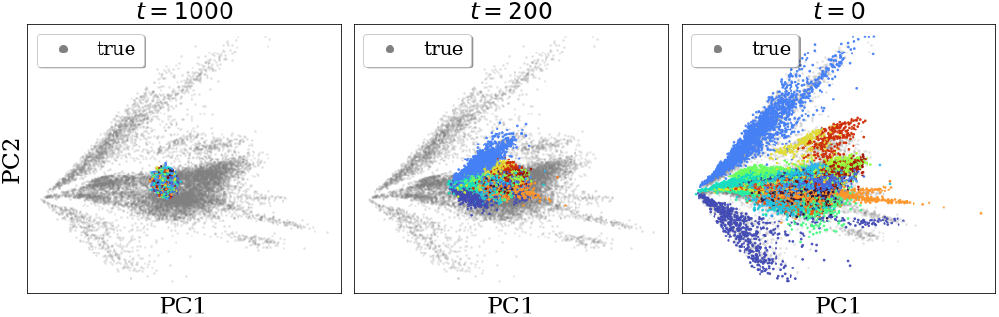
PCA visualization of the 1st and 2nd principal components throughout the generation steps *t* = 1000, …, 0 for GTEx data (L1000) and the DDIM-generated samples *x*_*t*_. Colors highlight the different tissue clusters.

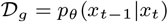

where *θ* denotes the parameters of the neural network optimized to estimate the loss function (below). In the original computer vision applications, the prevalent deep learning architecture is typically a U-NET (Ronneberger et al., 2015) with a stacking of autoencoders.

The loss function used to train the model is the KL divergence between the joint distributions of the forward and reverse Markov chains. In addition, the parameterization is simplified to yield a more smooth loss function. Along this line, the generative sampling produces a denoised sample *x*_*t−*1_ from *x*_*t*_ and *ϵ*_*θ*_:

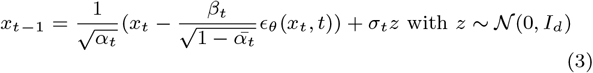

where *ϵ*_*θ*_ is the output of the trained neural network, computed from *x*_*t*_ and *t*. Simply put, the DM generative model is trained to directly predict the noise added on sample *x*_*t*_ at time step *t* during the forward process, of the same dimension as *x*_*t*_ (Fig. 1, panel C).

Eventually, the simplified loss function is the mean squared error:

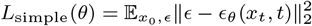

where *x*_*t*_ is computed from Eq. 2. The smoothness of the loss function leads to significantly more stable training compared to, e.g., the GAN min-max loss.

Despite the success of DDPMs, a notable challenge lies in the inference time required for generating samples. A generated sample is produced by: i) sampling *x*_*T*_ from the prior 𝒩 (0, *I*_*d*_); ii) *T* times, iteratively sampling *x*_*t−*1_ using *x*_*t*_ and *ϵ*_*θ*_(*x*_*t*_, *t*)). This process implies *T* passes through the neural network (as opposed to one pass in a VAE or a GAN). As many diffusion steps (*T ≈* 1, 000) are recommended to generate high-quality samples, time inference remains a major bottleneck for the practicality of such diffusion models.

#### Denoising Diffusion Implicit Models

Song *et al*. (2021) introduce Denoising Diffusion Implicit Models (DDIMs) to enforce a good trade-off between the quality of the generated sample and the computational cost by redefining the diffusion process as a non-Markovian process. The DDIM framework is based on the same objective function and backbone neural network as the DDPM one: it can be applied post-training on top of a previously trained DDPM model.

It proceeds by replacing the variance term *β*_*t*_ with *σ*_*t*_(*η, β*_*t*_) where *η* is a positive value that controls the stochasticity of the generation process: with the same notations as in Eq. 2,

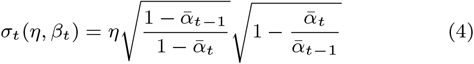

In the forward process (Eq. 1), *x*_*t*_ now depends on both *x*_*t−*1_ and the initial sample *x*_0_, making the forward process non-Markovian, though it preserves the closed-form property of the DDPM (Eq. 2). With the use of *σ*_*t*_(*η, β*_*t*_), the stochasticity of the forward process can be controlled. For *η* = 1, one falls back on the DDPM setting. For *η* = 0, the forward process is deterministic in the sense that *x*_*t−*1_ can be directly computed from *x*_0_ and *x*_*t*_. In such a case, the backward process is also (trained to be) deterministic: from a given input noise *x*_*T*_, the same generated sample *x*_0_ is always derived. This deterministic process entails nice properties, such as the ability to interpolate among generated samples by applying the backward process to the interpolation of their input noise samples.

Overall, the denoising step in Eq. 3 becomes:

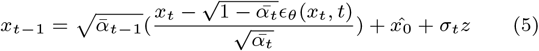

with 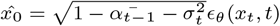 and 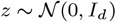.

The same loss function *L*_simple_ is used to optimize the model. The difference is that the denoising pass (Eq. 5) enables to skip intermediary steps to increase generation speed. When the process is deterministic, the resulting generated samples have the same high-level features and only differ in details, depending on the number of skipped steps.

According to the authors, DDIMs outperform DDPMs when the number of diffusion steps is lower than the initial *T* steps. The transition from DDPMs to DDIMs holds promise for addressing scalability challenges and extending the applicability of generative models across diverse domains.

### 2.2 Adapting DDIM to the high dimensional transcriptomics space

The proposed approach builds upon (Subramanian *et al*., 2017), which investigated leveraging redundancy and correlations in gene expression data to enhance sequencing efficiency and resource allocation. The authors introduce an ensemble comprising circa 1,000 landmark genes, referred to as the L1000 genes; they employ linear regression to infer the expression data of all other genes (circa 19,000) from the L1000 genes.

Subsequently, Chen *et al*. (2016) and Jeon *et al*. (2022) proposed neural learning approaches using the dimensionality reduction permitted by the L1000 genes to achieve gene expression inference via multilayer perceptrons (MLPs) architectures. Operating in the reduced L1000 space is shown to efficiently address the instability and memory consumption problems that can be encountered when training large deep generative models on such high-dimensional data.

The presented study proceeds by conducting the data generation phase in the L1000 landmark genes space noted *L*_*g*_. The generated samples are mapped to the remaining target genes space noted *T*_*g*_, using either a trained linear regression (LR) or a multilayer perceptron (MLP) method. Both methods are trained by minimizing the Mean Squared Error (MSE) between the true target genes and reconstructed genes (Eq. 6) on the validation set. The Mean Absolute Error (MAE) is additionally employed to facilitate a comprehensive comparison of the reconstruction performances, mitigating the influence of outliers. We refer to the complete transcriptome as the ensemble *L*_*g*_ ∪ *T*_*g*_.

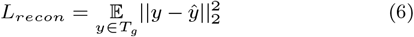

with *x* ∈ *L*_*g*_ the input in reduced dimension space, *y* ∈ *T*_*g*_ its representation in the target genes space, and *ŷ* = *ϕ*(*x*) its approximation by the reconstruction model *ϕ*. This reconstruction model is then used to map the synthetic landmark genes onto the target genes (Fig. 1, panel E).

## 3 Experimental setting

This Section describes the experimental settings for reliable model comparisons between a VAE, a WGAN-GP (Appendix J), and our DDIM. All final models are trained on a single NVIDIA A40 GPU with 48 GB of RAM. For statistical significance analysis, means and standard deviations are computed over five runs with the same setting.

### 3.1 Benchmark RNA-seq datasets

The presented experiments consider the Pan-Cancer Genome Atlas (TCGA) (Weinstein *et al*., 2013) and the Genotype-Tissue Expression (GTEx) (Lonsdale *et al*., 2013). TCGA data was retrieved using the RTCGA^1^ package in R (release date 2016-01-28), while the latest version (v8) of GTEx data (with TPM normalization) was obtained from the open-access portal.^2^ The preprocessing procedure, involving removal of duplicates, landmark and target genes IDs mapping, and standardization, can be found in our open-source code.^3^ The final TCGA dataset includes 6,499-1,625-1,625 train-validation-test samples of 978 landmark genes and 19,553 target genes to reconstruct. The GTEx dataset contains 9,796-2,448-5,000 train-validation-test samples of 974 landmark genes and 17,717 target genes. We consider 24 tissue types for TCGA and 26 for GTEx.

### 3.2 Data Quality Evaluation

Our data quality evaluation methodology follows the guidelines in Lacan *et al*. (2023), with unsupervised and supervised indicators presented below (see Appendix B for more details).

- Reverse Validation accuracy assesses the predictive capacity of generated data using an MLP trained on generated data and tested on the true test data unseen during training.
- Frechet Distance (FD) (Heusel *et al*., 2017) measures the similarity between true and generated data based on the last hidden layer of an MLP trained to discriminate tissue types.
- Precision and Recall (PR) (Kynkäänniemi *et al*., 2019) measure the distance between true and fake distributions considering the intrinsic dimensionality of their support.
- F1-score represents the harmonic mean between precision and recall measures.
- Adversarial Accuracy (AA)(Yale *et al*., 2020), assesses whether generated data balance accuracy and privacy in the context of sensitive data.
- Correlation Score (Viñas *et al*., 2021) compares the correlation matrices of true and generated data using Pearson correlation scores.

#### Unsupervised Indicators

Notably, Heusel *et al*. (2017) and Kynkäänniemi *et al*. (2019) highlight the sensitivity of FD and PR metrics to the number of considered samples, prompting compliance with the computation methodology from Lacan *et al*. (2023). These indicators were used to assess the quality of both the generated data and the reconstructed data, in addition to the MAE. A qualitative assessment was also performed using dimensionality reduction visualizations such as Uniform Manifold Approximation and Projection (UMAP) and Principal Components Analysis (PCA). As a reminder, we want to maximize all the similarity metrics, such as PR, F1-score, and correlations. On the contrary, we want to minimize the FD over the different configurations. Finally, the optimal value for the Adversarial Accuracy (AA) is 0.5 for a trade-off between privacy and performance.

#### Supervised Indicators

The baseline performance indicator based on the predictive accuracy is computed by training an MLP, optimized through a Bayesian optimization (Akiba *et al*., 2019) over 1,000 trials on the validation sets. We considered the multiclass classification task of the L1000 landmarks genes and the ensemble *L*_*g*_ ∪ *T*_*g*_) over tissue types. The reverse validation is based on the same best fixed MLP architecture trained on the generated data only.

### 3.3 Model Configuration

Deep generative models require a carefully selected training procedure due to their hyperparameter sensitivity. An ensemble of best hyperparameters was obtained for each generative model after maximizing the unsupervised F1 score (Section 3.2) through a Bayesian search of 100 trials. Embeddings of the tissue type covariates are used to condition all the generative models (Viñas et al. (2021)). Details of the VAE and WGAN-GP baseline models optimization can be found in Appendix A.

The DDIM is trained with many epochs (15,000) due to numerous diffusion steps (1,000) and a more expressive architecture. We adapted the architecture as a residual block of the same input and output size (Fig.1, panel C) for we could not use the typical U-NET model, unsuited for the characteristics of our tabular data. DMs also leverage the power of attention mechanisms and sophisticated class conditioning (e.g., classifier guidance in Ho and Salimans (2021)), which we did not implement in this first adaptation. We used Automatic Mixed Precision (Micikevicius *et al*., 2018) alongside a learning rate warmup strategy and big batch sizes (2,048) to keep an efficient training time. In addition to the residual block layers dimensions and the learning rate, we optimized the dropout rate, the variance (*β*_*t*_) scheduler (constant, linear, or quadratic), and the conditioning time steps (with or without sinusoidal embedding). Input data scaling was also investigated, and the scaling in [*−*1, 1] led to the best results. As stated by Song *et al*. (2021), this ensures that the neural network reverse process operates on consistently scaled inputs. The training time is 1 hour and 3 minutes for the DDIM on TCGA and 3 hours and 7 minutes for GTEx. The best retained hyperparameters can be found in Appendix A.

## 4 Results

This Section discusses the results obtained with the compared generative models: DMs, VAEs, and GANs, along with the performance indicators presented in Section 3.2.

### 4.1 Data generation in the reduced L1000 space

In Fig. 2, the GTEx DDIM-generated data are represented in the 2D plane defined from the PCA analysis at each diffusion step, where the colors correspond to the different tissues. The diffusion trajectories over the 1,000 diffusion steps show that synthetic data are initially grouped in the center of the image (*t* = 1, 000) and gradually spread across true data points (in grey) when *t* goes to 0, nicely grouping together the tissue-related clusters. Similar nicely defined clusters can be found in the TCGA UMAP (Appendix I).

The quantitative analysis is presented in Tables 1 and 2, reporting the unsupervised performance indicators on GTEx and TCGA, respectively. On GTEx, DDIM consistently ranks second (except in precision), while VAE or WGAN-GP ranks first, depending on the indicator. All the models exhibit strong correlations and precision, exceeding at least 98%, indicating that the generated data closely resemble true data. Regarding diversity (true clusters are all visited by generated data), VAE is significantly outperformed by WGAN-GP and DDIM (from circa 59% to circa 90%). VAE is also outperformed regarding the Frechet distance.

**Table 1.** Comparing the unsupervised indicators performance of the different generative models on GTEx. Statistically significantly best results are indicated in bold. *↑*: the higher, the better. *↓*: the lower, the better.

| Metric | VAE | WGAN-GP | DDIM |
| --- | --- | --- | --- |
| Corr. ↑ | 0.9814 ± 0.0008 | **0.9943** ± 0.0004 | 0.9893 ± 0.0033 |
| Prec. ↑ | **0.9944** ± 0.0006 | 0.9926 ± 0.001 | 0.9888 ± 0.0005 |
| Recall ↑ | 0.5928 ± 0.0021 | **0.9047** ± 0.0009 | 0.8923 ± 0.0028 |
| F1 ↑ | 0.7428 ± 0.001 | **0.9466** ± 0.001 | 0.9381 ± 0.0009 |
| AA | **0.5033** ± 0.0045 | 0.6047 ± 0.0027 | 0.4647 ± 0.0061 |
| FD ↓ | 4.0768 ± 0.1212 | **0.935** ± 0.0414 | 3.8931 ± 1.9 |

**Table 2.** Comparing the unsupervised indicators performance of the different generative models on TCGA. Statistically significantly best results are indicated in bold. *↑*: the higher, the better. *↓*: the lower, the better.

| Metric | VAE | WGAN-GP | DDIM |
| --- | --- | --- | --- |
| Corr. ↑ | 0.9826 ± 0.0004 | 0.9845 ± 0.0009 | **0.9918** ± 0.0002 |
| Prec. ↑ | 0.9976 ± 0.0003 | 0.9528 ± 0.0014 | **0.9999** ± 0.0001 |
| Recall ↑ | 0.7013 ± 0.0058 | **0.9116** ± 0.0023 | 0.7198 ± 0.0035 |
| F1 ↑ | 0.8236 ± 0.0006 | **0.9317** ± 0.0017 | 0.837 ± 0.0002 |
| AA | **0.4719** ± 0.0054 | 0.6904 ± 0.0053 | 0.4454 ± 0.0055 |
| FD ↓ | 0.6269 ± 0.0291 | 2.5603 ± 0.363 | **0.5181** ± 0.0339 |

In contrast, DDIM is better at generating TCGA data with high fidelity as the model ranks first regarding correlation, precision, and Frechet distance. However, looking at recall, WGAN-GP significantly outperforms the other models that fail to reach more than circa 70%/71% of diversity. A tentative interpretation is that both VAE and DDIM fail to capture sufficiently the heterogeneity of cancerous samples.

Regarding privacy, the adversarial accuracy (AA) is relatively stable around *~*0.5 for all models, except for WGAN-GP (0.6047 for GTEx, 0.6904 for TCGA), suggesting that it tends to generate data more easily distinguishable from the true dataset. The DDIM is second best after the VAE on both datasets.

The relationship between diversity and inference steps is shown in Table 3, comparing DDIM and DDPM (same trained model with *η* = 1) and for a number *T* of diffusion steps *{*50, 100, 1000*}* in the generation pass. The highest diversity is obtained with a DDPM using 1,000 diffusion steps for GTEx and TCGA; this result is consistent with a higher *η* leading to higher stochasticity in the generation process. In addition, a DDIM with 50 time steps yields a higher diversity (86.09%) than a DDPM with 100 steps (85.74%), thus dividing by half the inference time (from 28 sec. to 14 sec.). On TCGA, a DDPM with 100 steps reaches almost the same diversity (70.34%) as the best DDIM on 1,000 steps (71.70%), thus dividing by nine the inference time (from 127 sec. to 14 sec.). Skipping time steps in a DDIM framework can, therefore, efficiently decrease the generation cost while preserving good diversity (more in Appendix C).

**Table 3.** Comparative results in terms of recall (diversity) and the number of diffusion steps used in the generative process (the computational cost varies linearly with the number of diffusion steps). Statistically significantly better results are indicated in bold.

| Steps | GTEx |  |  | TCGA |  |  |
| --- | --- | --- | --- | --- | --- | --- |
|  | 50 | 100 | 1,000 | 50 | 100 | 1,000 |
| DDPM ($\eta = 1$) | 0.82 | 0.86 | **0.90** | 0.64 | **0.70** | **0.77** |
| DDIM ($\eta = 0$) | **0.77** | **0.86** | 0.89 | **0.65** | 0.68 | 0.72 |
| Inference time (s) | 14 | 28 | 268 | 6 | 14 | 127 |

Regarding computational resources, diffusion models require larger neural architectures than VAEs and WGANs-GP (Appendix H). Reaching the same realism-diversity performance on GTEx requires a DDIM architecture bigger by an order of magnitude than for WGAN-GP (228 million parameters vs 19 million). On TCGA, the best DDIM is almost three times bigger than the best WGAN-GP. The VAE, which hardly reaches the same performances as WGANs-GP or DDIMs, might also need a bigger neural architecture.

In counterpart to their being more computationally demanding, DMs are easier to train and deliver more stable results than VAEs or WGANs-GP. In our experiments on TCGA, for instance, circa 18% VAE runs (15% WGAN-GP runs) fail to converge.

### 4.2 Generation of complete gene expression profiles

In Appendix D and E, we first evaluated the linear regression and MLP reconstruction strategies (on true data). The MLP yields a better MAE, but both mappings lead to similar data quality regarding the unsupervised indicators. For the different generative model pipelines, we then considered both methods for synthetic L1000 genes reconstruction to *T*_*g*_ for a fair comparison.

Regarding the unsupervised indicators, each pipeline’s PR performances are reported in Fig. 3. On average, the MLP mapping yields good precision, sometimes at the expense of a small loss in recall compared to LR. On GTEx (Fig. 3, left), the best option is the WGAN-GP + mapping (similar to the performance obtained on L1000). On TCGA (Fig. 3, right), DDIM catches up with WGAN-GP, and both are on the precision-recall Pareto front. When observing the reverse validation accuracy in Fig.4, the main finding is that DM-generated L1000 data (DDPM and DDIM) is closest to reaching baseline classification performances (with true data) on both datasets. This shows DM-generated data can be used directly in the reduced landmark gene space as an alternative synthetic dataset.

**Fig. 3:**
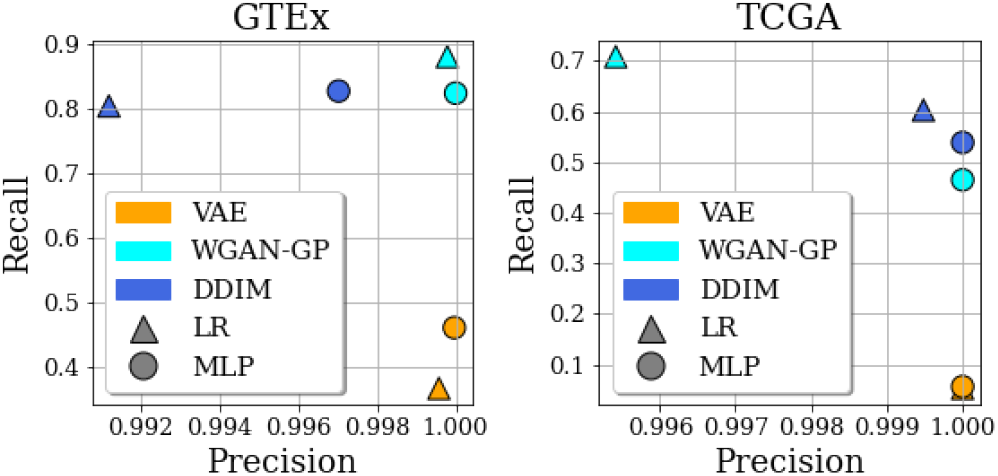
Precision/Recall trade-off according to the generative model, the reconstruction method and the dataset. Metrics are computed on the final sets of generated + reconstructed genes (*L*_*g*_ ∪ *T*_*g*_).

**Fig. 4:**
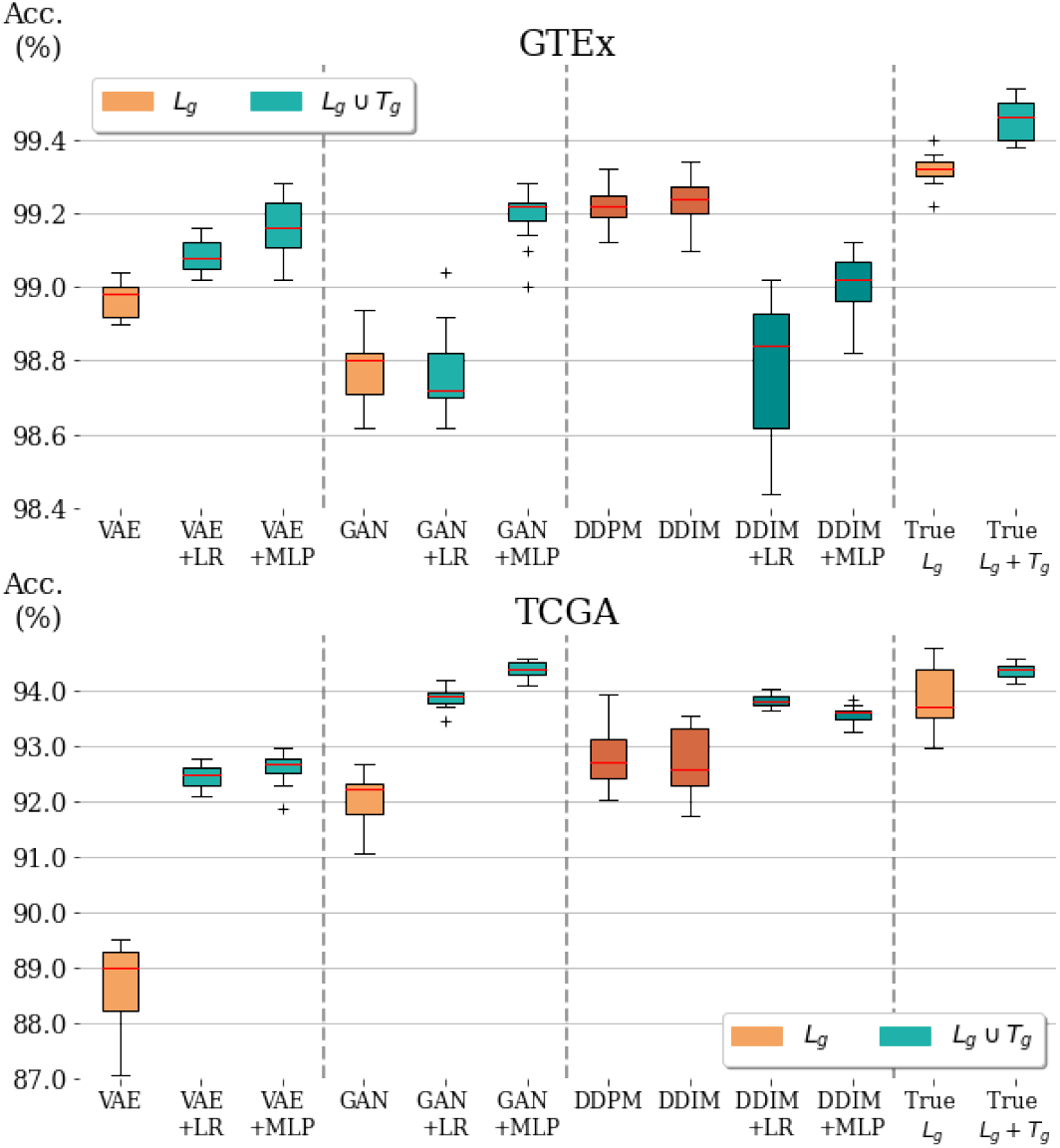
Predictive accuracy on tissue classification (supervised performance): on GTEx (top) and TCGA (bottom), compared with the baseline trained on the true data (rightmost). We differentiate between classification in reduced space *L*_*g*_ (orange), full gene space *L*_*g*_ ∪ *T*_*g*_ (blue) and DM-generated data in darker colors. All classifiers associated with a given dataset are trained on the same number of samples, share the same architecture.

Interestingly, all methods’ performances are generally improved when mapping the generated data to the entire gene space (except DDIM on GTEx, displaying a minimal gap in order of magnitude). For WGAN-GP, the predictive accuracy is better when considering the data generation in L1000, followed by a mapping on *T*_*g*_, compared to the data generation in *L*_*g*_ ∪ *T*_*g*_ directly (Appendix G). For VAE, we observe a similar accuracy pattern on GTEx. On a more heterogeneous dataset like TCGA, the WGAN-GP also benefits from the reconstruction method for the PR trade-off. It reaches 0.995/0.7 with LR mapping (Fig. 3) compared to 0.74/0.61 for the best WGAN-GP generating directly the entire transcriptome (Appendix F). In light of this comparative study between all the generative models, the heterogeneity and dimensionality of each dataset, the reconstruction method is a proper alternative to generating all the genes.

## 5. Conclusion

This paper presents two diffusion model frameworks, DDPM and DDIM, Moreover, it establishes that both can efficiently be used to achieve data generation for transcriptomics. These results, showing the first success of DMs application to transcriptomics to our best knowledge, rely on the ability to learn a generative model in a reduced 1,000-dimensional space and to map the generated data in the original high-dimensional transcriptome space.

The extensive comparative validation of these approaches on TCGA and GTEx datasets shows that DMs can be competitive with the alternative generative models, VAE and WGAN-GP. Regarding predictive accuracy in the reduced L1000 gene space, DM-generated data yields the closest performance to the baseline accuracy based on true data. On the positive side, DMs are easier to train in terms of stability and convergence of the models; on the negative side, the trained models are one order of magnitude larger than VAEs or GANs. Additionally, we observed that the mapping strategy to the full transcriptome helps all the generative models pipelines.

The perspectives of research opened by these approaches primarily concern the conditioning of the diffusion trajectories based on the patients covariates (e.g., age, gender, tissue type) and domain knowledge. Compared to GANs, such guidance of DMs is a substantial asset for the diversity aspect. Another interesting perspective is interpolating the generated data, e.g., to pave the representation space better and identify potential biomarkers.

## Footnotes

1 https://bioconductor.org/packages/RTCGA.

2 https://gtexportal.org/home/downloads/adult-gtex

3 https://forge.ibisc.univ-evry.fr/alacan/rna-diffusion.git

