## Appendix for "In Silico Generation of Gene Expression profiles using Diffusion Models"

### Supplementary materials

These supplementary materials are organized as follows:

- A. Best hyperparameters of all models considered
- B. Details about unsupervised data quality indicators
- C. Frechet distance evolution w.r.t. the number of diffusion steps (DDPM vs. DDIM)
- D. MAE of reconstruction methods
- E. Unsupervised data quality indicators obtained from reconstructed data
- F. Unsupervised data quality indicators obtained for data generated by VAEs and WGANs-GP trained directly on all landmark and target genes
- G. Reverse validation accuracy obtained for data generated by VAEs and WGANs-GP trained directly on all landmark and target genes
- H. Models complexity and training time
- G. UMAP visualization on TCGA

### Appendix A. Best hyper-parameters

In the interest of time and resources, we limited the search for the best hyper-parameters to the batch size, learning rate, optimizer, and hidden layers dimensions for the VAE and WGAN-GP over 1000 epochs. As numerical errors can arise with the VAE, it is useful to force the initialization of weights to minimal values ( 8.10-2). As regards the WGAN-GP, the discriminator requires five additional iterations than the generator, and each network needs its own learning rate so that none of them outperforms the other too quickly. The training time is 1 hour and 40 minutes for the best WGAN-GP on TCGA (batch size is 64), and 21 minutes on GTEx (batch size is 1024).

| | GTEx | TCGA |
|---:|---:|---:|
| batch_size | 256 | 2048 |
| hidden dim 1 (enc./dec.) | 8192 | 2048 |
| hidden dim 2 (enc./dec.) | 4096 | 8192 |
| hidden dim 3 (enc./dec.) | 4096 | 256 |
| hidden dim 4 (enc./dec.) | 2048 | 4096 |
| latent_dim | 128 | 128 |
| learning rate | 0.0008 | 0.001 |
| optimizer | Adam | Adam |

**Table 1.** Best hyperparameters for the VAEs trained on landmark genes.

| | GTEx | TCGA |
|---:|---:|---:|
| batch size | 1024 | 64 |
| hidden dim 1 (disc.) | 8192 | 4096 |
| hidden dim 1 (gen.) | 128 | 256 |
| hidden dim 2 (disc.) | 256 | 128 |
| hidden dim 2 (gen.) | 1024 | 1024 |
| hidden dim 3 (gen.) | 4096 | 8192 |
| learning rate (disc.) | 0.000582 | 0.000124 |
| learning rate (gen.) | 0.002205 | 0.000275 |
| optimizer | RMSprop | RMSprop |
| spectral normalization | False | False |

**Table 2.** Best hyperparameters for the WGANs-GP trained on landmark genes.

| | GTEx | TCGA |
|---|---|---|
| beta schedule | quadratic | quadratic |
| hidden dim | 8192 | 4096 |
| dropout | 0.1 | 0.1 |
| learning rate | 0.0005 | 0.001 |
| time sinusoidal embedding | False | False |

**Table 3.** Best hyperparameters for the DDIM trained on landmark genes.

| | GTEx | TCGA |
|---|---|---|
| batch size | 64 | 2048 |
| dropout | 0.3 | 0.5 |
| hidden dim 1 | 1024 | 256 |
| hidden dim 2 | 1024 | 16 |
| learning rate | 0.0001 | 0.004 |
| optimizer | Adam | Adam |

**Table 4.** Best hyperparameters for the baseline MLP trained on landmark genes.

| | GTEx | TCGA |
|---|---|---|
| batch size | 1024 | 64 |
| dropout | 0.5 | 0.5 |
| hidden dim 1 | 2048 | 1024 |
| hidden dim 2 | 8192 | 1024 |
| learning rate | 0.000516 | 0.012551 |
| optimizer | Adam | SGD |

**Table 5.** Best hyperparameters for the baseline MLP trained on all the genes (landmarks and targets).

### Appendix B. Unsupervised indicators details

We mainly used the range of indicators proposed by (Lacan et al., 2023).

#### Frechet Inception Distance (FD)

FD relies on the dimensionality reduction, achieved by mapping a sample $x$ onto its activations $y$ in some latent space e.g., the latent space of a discriminant neural network. The assumption is that true and generated data should have similar activation functions distributions. In computer vision, the FD is computed using the last pooling layer prior to the output classification of images in the Inception v3 pretrained network. Informally, the true and generated distributions are approximated as Gaussian distributions respectively noted $\mathcal{N}(\mu_t, \Sigma_t)$ and $\mathcal{N}(\mu_g, \Sigma_g)$. FD is the Wasserstein-2 distance between both Gaussian distributions (the lower, the better):

$$FD(,) = ||\mu_t - \mu_g||^2 + Tr(\Sigma_t + \Sigma_g - 2\sqrt{\Sigma_t.\Sigma_g}) \tag{1}$$

In computer vision, low FDs appear to be well correlated with higher quality generated images. FD is sensitive to the number of considered samples. We thus adapt the FD measure to transcriptomics by i/ considering all true training data samples; ii/ adjusting the embedding in the low-dimensional space considered to compute FD.

#### Precision and Recall (PR)

PR actually measure the distance between the true and fake distributions, accounting for the intrinsic dimensionality of their support. Formally, precision (respectively recall) is the probability of a sample from $D_g$ (resp. $D_t$) to fall within the support of $D_t$ (resp. $D_g$) ranging in $[0, 1]$. A high precision suggests that generated data are "close" to true data; a high recall suggests that no true data is "far" from generated data.

A manifold approximation is used to account for the intrinsic dimensionality of the supports. Formally, to each (true or generated) sample $x$ is associated a ball $B(x, r_k(x))$ with $r_k(x)$ the distance between $x$ and its $k$-th nearest neighbor in $R^{20,000}$. The coverage of these balls approximates the manifold $H$ containing $x$. The precision is thus defined as the percentage of generated samples falling in $H_t$

and conversely the recall is the percentage of true samples falling in $H_g$:

$$precision(x \in) = \begin{cases} 1, \text{iff } x \in H_t \\ 0, \text{otherwise} \end{cases} \quad recall(x \in) = \begin{cases} 1, \text{iff } x \in H_g \\ 0, \text{otherwise} \end{cases} \tag{2}$$

As the curse of dimensionality impacts the relevance of Euclidean distances, we set the number of neighbors $k$ to 10 (respectively 50) to enforce stable precision and recall measures on the L1000 genes (respectively all the transcriptome). Like FD, the precision and recall are sensitive to the number of samples, and we consider all true samples in estimating these indicators.

Another performance indicator is the F1-score which is the harmonic mean of the precision and recall measures. Analogously to the FD score, these metrics are computed on the true training set to benefit from a large amount of comparison samples.

#### Adversarial Accuracy (AA)

In the context of sensitive data, (Yale et al., 2020) developed the Adversarial Accuracy (AA) metric to assess whether the generated data are neither too far from true data (hindering their utility) nor too close (possibly entailing a breach of privacy). Formally, letting $d_{tt}(i)$ and $d_{tg}(i)$ respectively denote the min distance of the $i$-th true sample to another true (resp. generated) sample, and symmetrically, $d_{gt}(j)$ and $d_{gg}(j)$ denote the min distance of the $j$-th generated sample to another true (resp. generated) sample, it comes:

$$AA = \frac{1}{2}\left(\frac{1}{n}\sum_{i=1}^{n} 1(d_{tg}(i) > d_{tt}(i)) + \frac{1}{n}\sum_{i=1}^{n}(d_{gt}(i) > d_{gg}(i))\right) \tag{3}$$

AA can be understood as the accuracy of a 1-NN classifier discriminating true from generated data. When AA decreases to 0, this suggests that the generated data are too close copies of the true data (the generator overfits the training set). On the contrary, an AA close to 1 indicates that the generated data can easily be discriminated from the true data (the generator underfits the training set). Overall, a generative model with an AA around 0.5 should achieve a good trade-off between accuracy and privacy.

Similarly to precision and recall, AA is based on Euclidean distances in the data space; however, it is not sensitive to the number of nearest neighbors considered according to our experiments. The reported AA thus follows Eq. 3 is computed on a subset of true training samples due to time constraints.

#### Correlation score

While the above indicators globally assess the difference between true and generated data distributions, (Viñas et al., 2021) propose to compare the moments of both distributions. More precisely, the so-called Pearson correlation score is computed from the correlation matrices $M_t$ and $M_g$ associated with true and generated data, as follows:

$$\rho(M_t, M_g) = \sum_{i=1}^{d}\sum j = i + 1^d \frac{(M_{i,j;t} - M_{i,j;t})}{\sqrt{\sum_{k=1}^{d} M_{i,k;t}^2 \cdot \sum_{k=1}^{d} M_{i,k;g}^2}} \tag{4}$$

### Appendix C. Frechet distance w.r.t. the number of diffusion steps: DDPM vs. DDIM

| | GTEx | | | TCGA | | |
|---|---|---|---|---|---|---|
| steps | 50 | 100 | 1000 | 50 | 100 | 1000 |
| DDPM ($\eta = 1$) | **2.0396** | **1.7771** | **1.2114** | **0.2660** | **0.2140** | **0.1730** |
| DDIM ($\eta = 0$) | 4.0588 | 3.9795 | 4.0412 | 0.6572 | 0.6443 | 0.5154 |

**Table 6.** Mean Frechet distance evolution according to the number of diffusion steps used in inference. Best results for a given dataset and a given number of steps are highlighted in bold. The Frechet distance decreases with the number of steps.

### Appendix D. Mean Absolute Error of reconstruction methods

| | GTEx (MAE ↓) | TCGA (MAE ↓) |
|---|---|---|
| **LR** | 0.1979 ± 1e-5 | 0.375± 1e-5 |
| **MLP** | **0.1543 ± 5e-4** | **0.2696 ± 1e-3** |

**Table 7.** Mean absolute reconstruction error on test dataset of the best linear regression (LR) and deep learning (MLP) methods. Best results in bold.

### Appendix E. Unsupervised indicators of reconstructed data

| | LR | MLP |
|---|---|---|
| **correlation ↑** | **0.9934±0.0002** | 0.9931±0.0004 |
| **precision ↑** | **1.0±1e-6** | **1.0±1e-6** |
| **recall ↑** | 0.976±1e-6 | **0.9855±0.0005** |
| **f1 ↑** | 0.9879±1e-6 | **0.9927±1e-6** |
| **AA** | 0.0954±0.0032 | **0.1536±0.0027** |

**Table 8.** Unsupervised indicators performance on the GTEx dataset w.r.t both reconstruction methods. Best results in bold.

| | LR | MLP |
|---|---|---|
| **correlation ↑** | **0.9779±0.0008** | 0.9748±0.0018 |
| **precision ↑** | **1.0±0.0** | **1.0±0.0** |
| **recall ↑** | **0.954±0.0** | 0.9302±0.0249 |
| **f1 ↑** | **0.9765±0.0** | 0.9638±0.0 |
| **AA** | 0.1598±0.0064 | **0.293±0.0271** |

**Table 9.** Unsupervised indicators performance on the TCGA dataset w.r.t both reconstruction methods. Best results in bold.

### Appendix F. Unsupervised indicators of data generated by the best VAE and WGAN-GP on all landmark and target genes

| | VAE | WGAN-GP |
|---|---|---|
| **correlation ↑** | **0.9922±0.0003** | 0.9894±0.0003 |
| **precision ↑** | **0.9993±0.0003** | 0.9973±0.0003 |
| **recall ↑** | 0.9274±0.0024 | **0.9381±0.0018** |
| **f1 ↑** | 0.962±0.0005 | **0.9668±0.0005** |
| **AA** | 0.4789±0.0039 | **0.5847±0.005** |

**Table 10.** Unsupervised indicators performance on the GTEx dataset w.r.t the best VAE and WGAN-GP trained on both landmark and target genes. Best results in bold.

| | VAE | WGAN-GP |
|---|---|---|
| **correlation ↑** | **0.9708±0.0006** | 0.7312±0.032 |
| **precision ↑** | **1.0±1e-6** | 0.7457±0.0036 |
| **recall ↑** | **0.7918±0.0036** | 0.615±0.0011 |
| **f1 ↑** | **0.8838±0.0** | 0.6741±0.0017 |
| **AA** | **0.4699±0.0022** | 0.8243±0.005 |

**Table 11.** Unsupervised indicators performance on the TCGA dataset w.r.t the best VAE and WGAN-GP trained on both landmark and target genes. Best results in bold.

### Appendix G. Reverse validation accuracy for the best VAE and WGAN-GP trained on all landmark and target genes

| | GTEx | TCGA |
|---|---|---|
| VAE | 0.2674±0.1045 | 0.9449±0.0027 |
| WGAN-GP | 0.9916±0.0007 | 0.937±0.002 |

**Table 12.** Reverse validation on true tissue types: test accuracy of a MLP trained only on a generated dataset of all landmark and target genes. All classifiers are trained on the same number of samples and share the same architecture.

### Appendix H. Models complexity and training time

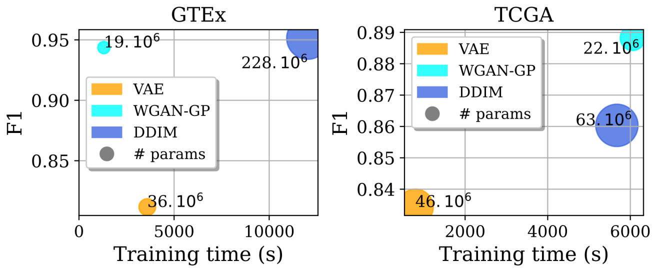


Fig. 1: Generative models complexity: number of parameters of the best generative models w.r.t. training time and unsupervised F1 indicator.

### Appendix I. UMAP visualization

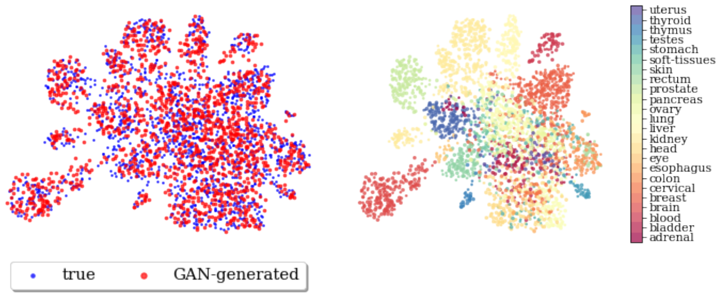


Fig. 2: TCGA dataset (L1000): true and GAN-generated data using UMAP visualization. Left: contrasting true and generated samples. Right: contrasting the 24 tissue clusters (in colors).

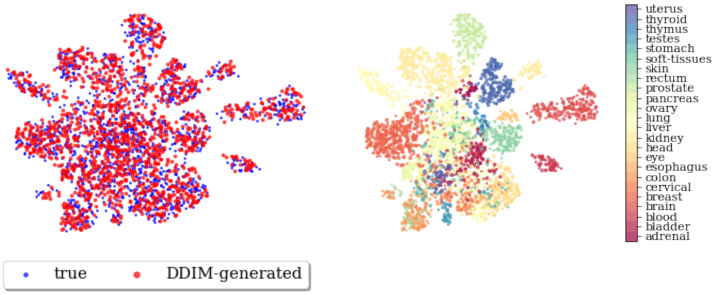


Fig. 3: TCGA dataset (L1000): true and DDIM-generated data using UMAP visualization. Left: contrasting true and generated samples. Right: contrasting the 24 tissue clusters (in colors).

### Appendix J. Variational Autoencoders and Generative Adversarial Networks

This Section briefly reminds the main generative models used for gene expression so far: Variational Encoders and Generative Adversarial Models.

**Notations.** In the following, $\mathcal{D}_t$ denotes the (unknown) true data distribution, and $\mathcal{D}_g$ the learned distribution parameterized from the model parameters $\theta$. Deep generative models proceed by learning a distribution $\mathcal{D}_g = p_\theta(x)$ that approximates the target distribution $\mathcal{D}_t$, supporting a sampling mechanism.

#### Variational Autoencoders

Variational Autoencoders extend the famed auto-encoder architecture as follows. The *encoder* maps input $x \in \mathbb{R}^D$ into the parameters of a distribution noted $Q(z|x)$ in the latent space. The *decoder* samples some $z \in \mathbb{R}^d$ after $Q(z|x)$, and maps $z$ onto a sample $x'$ in $\mathbb{R}^D$ generated from $P(x|z)$. The VAE learning criterion (Eq. 5) involves two terms; the former one is the reconstruction error (the MSE between $x$ and $x'$ in the case where $P(x|z)$ is a Gaussian distribution, up to a multiplicative constant). The latter is the Kullback-Leibler divergence between the image distribution $Q(z|x)$ and the target distribution in latent space (usually a Gaussian distribution).

$$\mathop{\mathbb{E}}_{z \sim Q(z|x)} [\log(P(x|z)) - \mathrm{KL}(Q(z|x)||p_\theta(z))] \tag{5}$$

#### Generative Adversarial Networks

A Generative Adversarial Network (GAN) comprises two components, a generator $G$ and a discriminator $D$ interacting along a zero-sum game. From the true data samples $x \in \mathcal{D}_t$, the generator $G$ learns a distribution $\mathcal{D}_g$ and uses it to generate synthetic data. The discriminator $D$ aims to discriminate between the true and the generated data, while the generator seeks to fool the discriminator. Informally, the generator succeeds when the discriminator fails to distinguish between $\mathcal{D}_t$ and $\mathcal{D}_g$. On one hand, GANs capture complex data distributions more easily than VAEs (e.g., when dealing with multimodal distributions). On the other hand, GANs face some challenges in optimizing the min-max learning criterion efficiently. The introduction of Wasserstein GANs (WGANs), particularly the WGAN-GP, addresses these challenges and offers enhanced stability. It mitigates issues arising from non-overlapping distributions by considering their Wasserstein distance via gradient penalties:

$$\begin{aligned} \min_G \max_D \quad & L(D,G) = \mathbb{E}_{x \sim \mathcal{D}_t}[D(x|c)] - \mathbb{E}_{z \sim P_z}[D(G(z|c))] \\ & + \lambda \mathbb{E}_{\tilde{x} \sim \mathcal{D}_g}[(\|\nabla_{\tilde{x}} D(\tilde{x})\|_2 - 1)^2] \end{aligned} \tag{6}$$

The $\lambda$ penalty weight enforces $D$ as a 1-Lipschitz function, and $\tilde{x}$ is the linearly interpolated data between true $x$ and generated $\hat{x}$ conditioned by context $c$.
